# Phage cocktail strategies for the suppression of a pathogen in a cross-feeding coculture

**DOI:** 10.1101/2020.06.08.140301

**Authors:** Lisa Fazzino, Jeremy Anisman, Jeremy M. Chacón, William R. Harcombe

**Affiliations:** Department of Microbiology and Immunology, University of Minnesota, Minneapolis, MN, USA; BioTechnology Institute, University of Minnesota, Saint Paul, MN, USA; College of Continuing and Professional Studies, University of Minnesota, Minneapolis, MN, USA; Department of Diagnostic and Biological Sciences, School of Dentistry, University of Minnesota, Minneapolis, MN, USA; Department of Evolution, and Behavior, University of Minnesota, Saint Paul, MN, USA

## Abstract

Cocktail combinations of bacteria-infecting viruses (bacteriophage), can suppress pathogenic bacterial growth. However, predicting how phage cocktails influence microbial communities with complex ecological interactions, specifically cross-feeding interactions in which bacteria exchange nutrients, remains challenging. Here, we used experiments and mathematical simulations to determine how to best suppress a model pathogen, *E. coli*, when obligately cross-feeding with *S. enterica*. We tested whether the duration of pathogen suppression caused by a two-lytic phage cocktail was maximized when both phage targeted *E. coli*, or when one phage targeted *E. coli* and the other its cross-feeding partner, *S. enterica*. Experimentally, we observed that cocktails targeting both cross-feeders suppressed *E. coli* growth longer than cocktails targeting only *E. coli*. Two non-mutually-exclusive mechanisms could explain these results: 1) we found that treatment with two *E. coli* phage led to the evolution of a mucoid phenotype that provided cross-resistance against both phage, and 2) *S. enterica* set the growth rate of the co-culture, and therefore targeting *S. enterica* had a stronger effect on pathogen suppression. Simulations suggested that cross-resistance and the relative growth rates of cross-feeders modulated the duration of *E. coli* suppression. More broadly, we describe a novel bacteriophage cocktail strategy for pathogens that cross-feed.

**Originality-Significance Statement:** Cross-feeding, or exchanging nutrients among bacteria, is a type of ecological interaction found in many important microbial communities. Furthermore, cross-feeding interactions are found to play a role in some infections, and research into treating infections with combinations of bacteriophage in ‘cocktails’ is growing. Here, we used a combination of mathematical modelling and wet-lab experiments to optimize suppression of a model pathogen with a bacteriophage cocktail in a synthetic cross-feeding bacterial coculture. A key finding was that a physiological parameter – growth rate – of the bacteria was important to consider when choosing the most effective cocktail formulation. This work is novel because it highlights an unexpected multispecies-targeting strategy for designing phage cocktails for cross-feeding pathogens and has relevance to many ecological systems ranging from human health to agriculture. We demonstrate how leveraging knowledge of a pathogen’s ecological interaction has the potential to improve precision medicine and management of microbial systems.

## Introduction

Phage have been used to treat pathogenic bacteria in human health, agriculture, and the food industry. Phage therapy and biocontrol often use multiple phage simultaneously in ‘cocktails’ to suppress pathogen growth (Pires *et al.*, 2017; Culot *et al.*, 2019; Gordillo Altamirano and Barr, 2019; Kakasis and Panitsa, 2019; Mahony *et al.*, 2020). Attacking a bacterial population with multiple phage can reduce the rate at which phage resistance evolves (Filippov *et al.*, 2011; Gu *et al.*, 2019; Ramírez *et al.*, 2020). However, we understand less about how treatment outcomes are affected by complex interactions among bacteria in a microbial community (Fazzino *et al.*, 2020). One bacterial interaction of particular interest is cross-feeding, in which metabolites secreted by one bacterium are used as a nutrient source by another. This is a common interaction in natural systems (Schink, 2002; D’Souza *et al.*, 2014; Mee *et al.*, 2014; Zelezniak *et al.*, 2015; Adamowicz *et al.*, 2018). Understanding how complex ecological interactions involving pathogens affect phage treatment outcomes will be critical for designing effective therapies. Here, we explore how two important factors - the potential for cross-resistance evolution and relative bacterial growth rates - interact with targeting strategies to suppress growth of a focal pathogen cross-feeding in an engineered coculture.

Experiments using cocultures with well-defined interactions have helped elucidate a range of responses to phage infection which may be leveraged for phage therapy. For example, adding non-host bacteria that compete with phage hosts for nutrients limits phage resistance evolution, thereby magnifying the efficacy of the phage (Harcombe and Bull, 2005; Brockhurst *et al.*, 2006; Wang *et al.*, 2017; Yu *et al.*, 2017; Testa *et al.*, 2019). Microbes also can engage in cooperative mutualistic interactions, where bacteria depend on others to cross-feed nutrients (Schink, 2002; Mee *et al.*, 2014; Zelezniak *et al.*, 2015; Souza *et al.*, 2016; Adamowicz *et al.*, 2018). Targeting one species in a cross-feeding mutualism can reduce the population of both mutualists, leading to the hypothesis that phage therapies could target a pathogen’s mutualists (Fazzino *et al.*, 2020). However, it is unknown how cocktails should be assembled to maximize pathogen suppression in a community.

If pathogenic bacteria cross-feed with other community members, then we can consider novel strategies of phage cocktail design that also target the nonpathogenic cross-feeding partner. Phage cocktails classically contain multiple phage that target a focal species to better limit the growth of a pathogen while also decreasing the rate of resistance evolution (Chan and Abedon, 2012; Wang *et al.*, 2017; Betts *et al.*, 2018). However, cross-resistance can evolve during treatment with classic pathogen-targeting cocktails when a single mutation blocks infection to multiple phage (Cairns and Payne, 2008; Wei *et al.*, 2011; Kortright *et al.*, 2019). We hypothesize that if a focal pathogen is engaged in an obligate mutualism, including a phage that targets the cross-feeding nonpathogen will increase suppression of the pathogen. It has been shown that off-target inhibition of cross-feeding partners can inhibit focal bacterial strains (Shou *et al.*, 2007; Adamowicz *et al.*, 2018). Combining phage that target the pathogen and its cross-feeding partner in a ‘multispecies-targeting’ cocktail would require the coculture to evolve two resistance mutations - one in each cross-feeding partner - to continue growing. Here, we hypothesize that this novel cocktail strategy that targets pathogens and cross-feeding nonpathogens will eliminate cross-resistance evolution and lengthen pathogen suppression.

In this study, we test the efficacy of phage cocktail treatment strategies to suppress a model pathogen obligately cross-feeding in a synthetic coculture. We performed wet-lab experiments with an engineered obligate cross-feeding coculture of an *Escherichia coli* methionine auxotroph that provides carbon to a methionine-secreting *Salmonella enterica* (Harcombe, 2010). Here, *E. coli* is the model pathogen to be suppressed. We introduced all pairwise combinations of *E. coli*-specific T7 and/or P1*vir* lytic phage, and the *S. enterica*-specific P22*vir* lytic phage (Fig. 1a). We then compared ‘pathogen-targeting cocktails’ with ‘multispecies-targeting cocktails’. We hypothesized and observed that targeting both cross-feeding partners was more effective at suppressing *E. coli* than targeting only *E. coli* with cocktails in our wet-lab experiments. We combined wet-lab experiments and mathematical modeling to uncover two reasons for this. First, as anticipated, we found evidence that *E. coli* evolved cross-resistance in the pathogen-targeting cocktail treatment. Second, the multispecies-targeting cocktail inhibited the slowest-growing cross-feeding partner, *S. enterica*, which limited how fast the coculture recovered from phage treatments. In fact, treatment with a single phage infecting *S. enterica* was as effective as targeting both cross-feeding partners in wet-lab experiments and simulations. Ultimately, our study highlights a novel strategy for designing phage cocktails that suppress cross-feeding pathogens.

**Figure 1.**
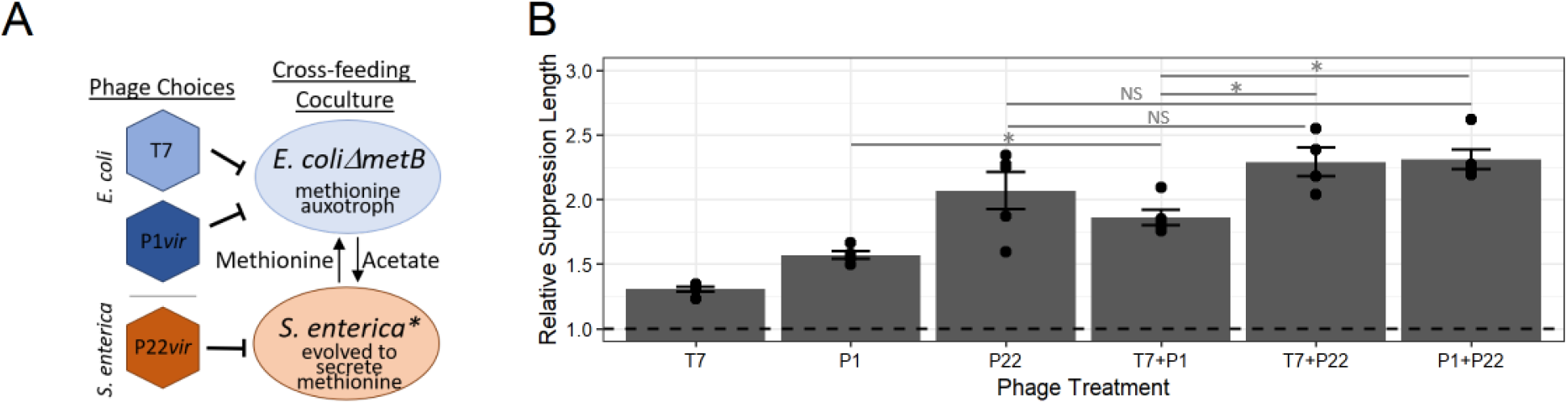
Phage cocktail and phage component suppression of cross-feeding microbial community. **A)** Schematic representation of the wet-lab engineered cross-feeding bacterial system with phage strains. *E. coli* = methionine auxotroph with cyan fluorescent protein, *S. enterica* = methionine secreter with yellow fluorescent protein. **B)** Relative *E. coli* suppression lengths of single and cocktail phage treatments standardized to the no phage control. Suppression length was calculated using 95% maximum cyan fluorescent protein measurement. Permutation statistical tests determined significance. p>0.1 (NS), p<0.05 (*). Exact p-values are in the text. Bars represent means ± SE (n = 4-5)

## Results

We wondered whether the pathogen, *E. coli*, would be suppressed for longer by a phage cocktail combining two *E. coli*-targeting phage (‘pathogen-targeting cocktail’) or combining an *E. coli*-targeting phage with a *S. enterica*-targeting phage (‘multispecies-targeting cocktail’). We grew control cocultures without phage (‘phage-free’), and treatment cocultures with combinations of T7 and P1*vir* as *E. coli*-targeting phage and P22*vir* as a *S. enterica*-targeting phage. The growth of each strain was tracked with unique fluorescent proteins (see Experimental Procedures for details). We predicted that the multispecies-targeting cocktail would provide the longest *E. coli* suppression because cross-resistance would not be possible and two mutations in separate bacterial species would need to evolve for the coculture to grow.

We quantified duration of *E. coli* suppression across phage treatments. To do this, we measured the amount of time required for a fluorescent protein in *E. coli* to reach 95% maximum intensity, which we refer to as time to maximum density. We calculated the relative suppression duration caused by each phage treatment as a fold-change relative to the phage-free cocultures, which required 34.4 hours to reach maximum density (Table S1). As anticipated, all phage treatments increased *E. coli* suppression duration (Fig 1b, Table S1 for absolute values). Notably, both of the multispecies-targeting cocktails delayed *E. coli* growth longer than the pathogen-targeting cocktail (p<0.02 for T7+P22*vir* and P1+P22*vir*), but were not significantly different from each other (p=0.43). Yet, the single phage treatment with the *S. enterica*-targeting phage P22*vir* suppressed *E. coli* equally as long as any of the cocktail treatments (p>0.17 for any cocktail) (Fig 1b, Table S1).

Two possible, but not mutually exclusive, reasons that targeting *S. enterica* suppressed *E. coli* growth longest are 1) that *E. coli* evolved cross-resistance to both phage in the pathogen-targeting cocktail, reducing its efficacy and/or 2) that *S. enterica* sets coculture growth rate, and that targeting it maximizes suppression of both species including *E. coli*.

We hypothesized that cross-resistance may be one reason why the pathogen-targeting cocktail was less effective than the multispecies-targeting cocktails. Multiple studies have reported that phage cocktails suppress focal bacteria less than expected given single phage treatments, suggesting that evolution of cross-resistance may be common (Cairns and Payne, 2008; Wei *et al.*, 2011; Kortright *et al.*, 2019). To determine if cross-resistance evolved, we measured phage resistance of *E. coli* isolates from each coculture with cross-streak assays. As expected, *E. coli* isolates from phage-free controls were sensitive to both *E. coli* phage (Table 1). Additionally, *E. coli* clones treated with a single phage were resistant to that phage, but remained sensitive to phage with which they had not been treated. Half of the *E. coli* clones from pathogen-targeting cocktail treatments evolved resistance to both *E. coli* phage, suggesting that cross-resistance may have evolved in some replicate cocultures. We also observed that all *E. coli* isolates from the pathogen-targeting cocktail treatments evolved mucoid phenotypes, which has previously been shown to cause cross-resistance by a single mutation in various genes involved in lipopolysaccharide production (Radke and Siegel, 1971; Skurray *et al.*, 1974; Mizoguchi *et al.*, 2003; Scanlan and Buckling, 2012).

**Table 1.** Resistance profiles and mucoid phenotypes of *E. coli* isolates to *E. coli*-specific phage.

| Treatment | T7 Resistant<sup>a</sup> / Total Reps | P1vir Resistant<sup>a</sup> / Total Reps | Mucoid / Total Reps |
| --- | --- | --- | --- |
| No Phage | 0/5 | 0/5 | 0/5 |
| T7<sup>b</sup> | 4/4 | 0/4 | 0/4 |
| P1vir | 0/5 | 5/5 | 0/5 |
| T7 + P1vir<sup>b</sup> | 2/4 | 4/4 | 4/4 |

<sup>a</sup> A representative isolate per treatment replicate was cross-streaked against the indicated phage.
<sup>b</sup> One repeat each of a T7-only treated community and a T7+P1vir-treated community had no detectable *E. coli* at the end of growth and were omitted for phenotyping.

Additionally, we wondered whether the observed resistances to multiple phage were caused by two independent mutations or a single mutation conferring cross-resistance. We predicted how common resistance to both *E. coli* phage would be in populations unexposed to phage (i.e. standing variation of resistance). If resistance to the two *E. coli* phage required different mutations, then the frequency of resistance to both phage is the likelihood that each resistance mutation was acquired individually (*f*_*cross-resistance*_ = *f*_mut1_ × *f*_mut2_). To quantify standing variation of resistance, we compared the number of resistant colonies on a phage-covered plate with the number of colonies on a phage-free plate for each ancestral bacteria (Luria and Delbrück, 1943). The frequency of resistance to both *E. coli* phage in the ancestral *E. coli* population was ~100-fold larger than predicted if resistance to both phage required two independent mutations (Fig 2 - T7+P1vir and red asterisk). These data suggest that the evolution of cross-resistance may be one reason the pathogen-targeting cocktail was less efficacious than the multispecies-targeting cocktail.

**Figure 2.**
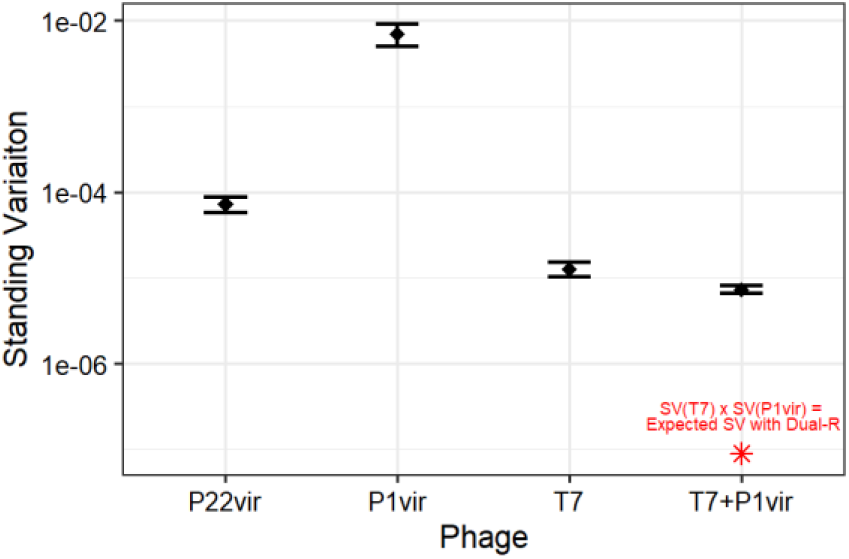
Resistance to phage standing genetic variation of ancestral bacterial species previously unexposed to phage. Standing variation frequencies are the number of bacterial colonies on plates with phage standardized to the number of colonies on plates without phage (black diamonds = means ± SE (n = 3)). Expected standing variation if dual-resistance occurred was calculated by multiplying the frequency of standing variation of T7 and P1*vir* (red asterisk).

Cross-resistance cannot explain another result: that a single phage targeting *S. enterica* is just as efficacious at increasing suppression of *E. coli* as any of the best cocktail treatments. A hypothesis which could explain this result is that *S. enterica*’s ability to recover from phage infection in the multispecies-targeting cocktail treatment set the coculture growth response. We tested two possible ways that *S. enterica* could act as a response-setting species: 1) that resistance to *S. enterica* phage was the least likely to evolve (i.e. smallest standing variation of phage resistance), or 2) that low *S. enterica* density caused by phage-mediated lysis delayed *E. coli* growth more than low *E. coli* density delayed *S. enterica* growth (i.e. *S. enterica* is the rate-limiting member of the co-culture).

To determine whether resistance to *S. enterica* phage was less likely to evolve than resistance to *E. coli* phage, we measured the standing variation in phage-free cultures of ancestral bacterial populations, or the frequency of resistance without exposure to phage. If the ability to evolve resistance was the determining factor of cocktail efficacy, then resistance to the *S. enterica* phage P22*vir* should have the lowest frequency of standing variation for resistance. Resistance to P22*vir* was more common than resistance to T7 (p=0.050, Fig 2) and less common than resistance to P1*vir* (p = 0.047, Fig 2), suggesting that resistance to T7, not P22*vir*, was hardest to evolve. If the ease of evolving resistance was the sole factor determining cocktail efficacy, then our results suggest that the longest suppression of *E. coli* should be caused by any treatment containing T7 phage. Yet, we observed that treatments including *S. enterica* phage P22*vir* suppressed *E. coli* growth the longest.

Alternatively, physiological differences between *S. enterica* and *E. coli* could cause a different co-culture response to phage. Other studies have illustrated asymmetrical responses of cross-feeding systems to perturbations caused by differences in growth rates or production rates of cross-fed nutrients (Shou *et al.*, 2007; La Sarre *et al.*, 2017; McCully *et al.*, 2017). Here, we examined the influence of physiology on co-culture rebound after a population size reduction by manipulating starting coculture frequencies without phage. This manipulation isolated the impact of phage-mediated population size reduction from the impact of phage replication and/or resistance evolution. We started cocultures with either *E. coli* or *S. enterica* at 0.01% instead of 50% of the coculture population and tracked *E. coli* growth as before. We found that reducing the density of *S. enterica* lengthened the time to *E. coli* maximum density more than reducing the starting density of *E. coli* (p <0.01, Fig 3).

**Figure 3.**
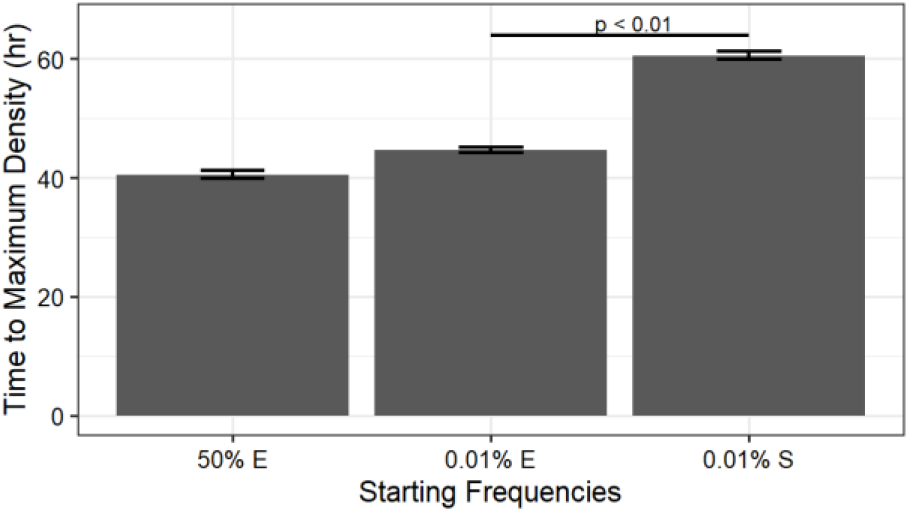
Time to maximum *E. coli* density when bacterial starting frequencies were altered in phage-free cocultures. Cocultures were grown as before with different initial starting densities of *E. coli* (E) or *S. enterica* (S). Statistics performed with permutation analysis. Means ± SE (n = 3).

Given that cross-resistance and phage-target identity both influenced the cocktail efficacies, we wondered whether one was more important to consider when predicting outcomes of cocktail treatment. We used a mathematical model to investigate this question. To start, we modified a resource-explicit model of the coculture that simulated the abundance of bacteria, phage, and resources through time (Fig 4a)(Fazzino *et al.*, 2020). We used model parameters informed by literature values and used wet-lab experiments to measure maximum growth rates (Table S2, Fig 4b), and confirmed that the model accurately simulated the growth dynamics of the phage-free coculture (Fig 4c)(Fazzino *et al.*, 2020). Phage-resistant bacteria were seeded in at low frequencies to approximate standing variation for phage resistance. We included two different phage resistance mechanisms and then manipulated phage target identity. Phage resistant mutants were either cross-resistant (resistant to two phage via one mutation) or dual-resistant (resistant to two phage via two independent mutations). The only difference between modeling resistance mechanisms was the doubly-resistant mutants’ starting frequencies. For cross-resistance, we seeded in mutants resistant to both phage at a frequency equal to the sum of single resistant mutant frequencies because resistance to either phage confers resistance to the other. For dual-resistance, doubly-resistant mutants were seeded in at a frequency equal to the product of single resistant mutant frequencies to approximate the likelihood that two independent mutations evolved by chance (Table S3). With this model, we simulated treatment with single phage and cocktail treatments. As expected, cross-resistance to two *E. coli* phage decreased the time to maximum *E. coli* density in pathogen-targeting cocktail treatments and the multispecies-targeting cocktails suppressed *E. coli* better than the pathogen-targeting cocktail when cross-resistance evolved (Fig 4d - dark bars). Furthermore, the relative efficacy of the cocktails depended on the resistance type. When we simulated cross-resistance, the multispecies-targeting cocktail was most effective (Fig 4d - dark bars); however, when we simulated dual-resistance the pathogen-targeting cocktail was most effective (Fig 4d - light bars). These simulations suggest that for our experimental coculture, the evolved resistance type determines which cocktail treatment is most effective. Note though, for both simulated resistance types, the single P22*vir* phage treatments targeting *S. enterica* suppressed communities equally as well as the multispecies-targeting cocktail, suggesting that phage-target identity also contributes to treatment efficacy.

**Figure 4.**
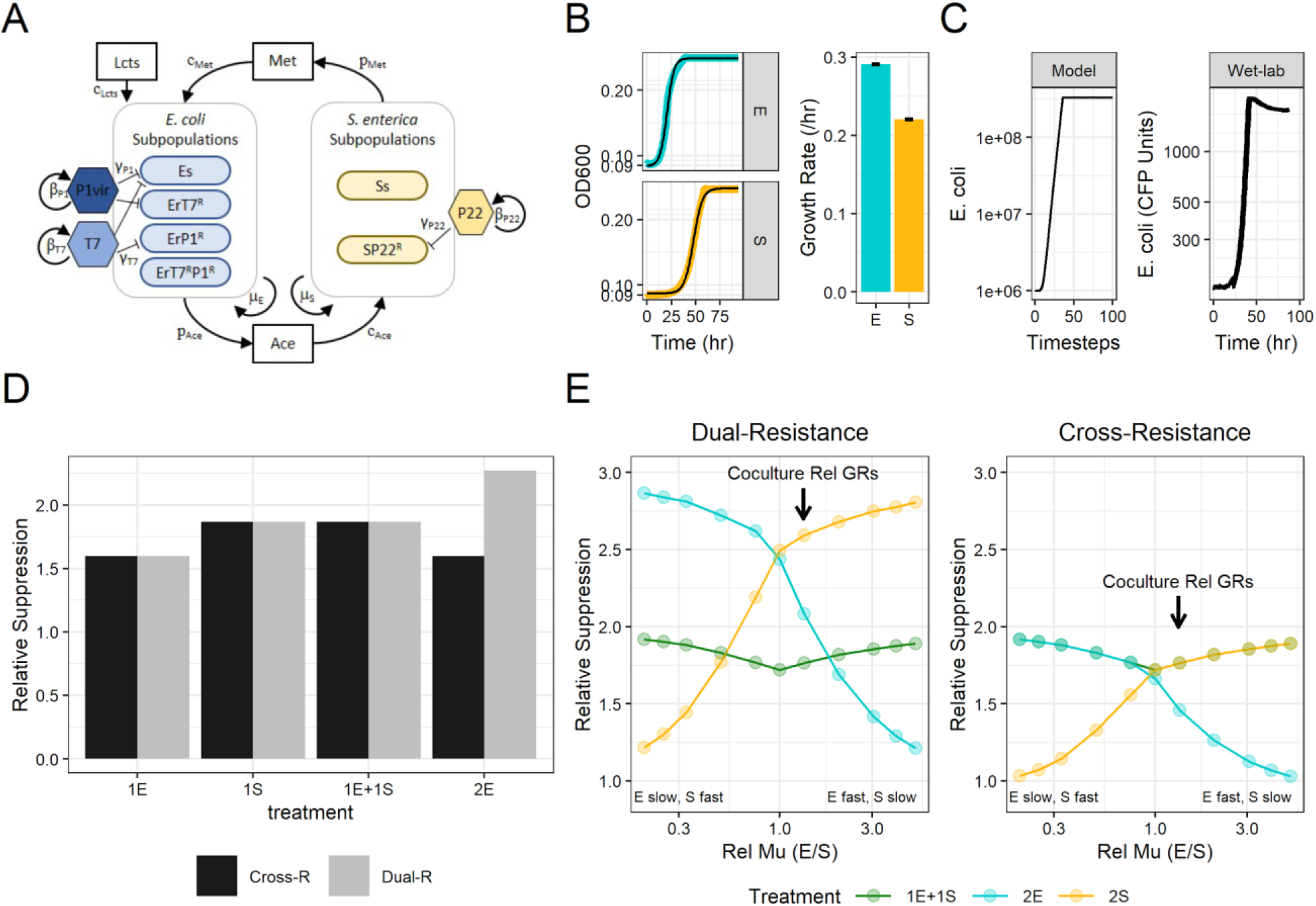
Simulations of coculture growth with phage treatments. **A)** Schematic showing cross-feeding interactions between *E. coli* (E) and *S. enterica* (S) subpopulations. Simulated bacterial subpopulations are listed in species boxes and allowed tracking sensitive (Xs) and phage-resistant (X^R^) populations of *E. coli* (E) or *S. enterica* (S). Key tracked metabolites are in boxes. Arrows show direction of interactions. Key model parameters are next to associated arrows: μ_x_ = maximum growth rate of species X; p_m_ = production rate of metabolite; c_m_ = consumption rate of metabolite m; β_v_= burst size of phage V; γ_v_ = adsorption rate of phage V. See Table S2 for details. **B)** Parametrizing bacterial growth rates from wet-lab data. The left panel are representative OD600 growth curves of *E. coli* (blue) and *S. enterica* (yellow) monocultures overlaid with Baranyi growth fits (black lines). The right panel shows calculated growth rates for each species. Bars are means ± SE (n = 5). **C)** Comparison of *E. coli*-specific phage-free coculture growth curves from the model and wet-lab experiments. Y-axis of the model growth curve is the total simulated *E. coli* biomass and the y-axis of the wet-lab growth curve is measured with CFP fluorescence units. **D)** Relative suppression (time to maximum *E. coli* density relative to phage-free simulations) of either cross-resistance (Cross-R) or dual-resistance (Dual-R) simulations with experimentally determined growth rates. Simulating resistance mechanisms used different starting densities of phage-resistant subpopulations (see text and experimental procedures for details). **E)** Simulation of relative suppression while modulating relative bacterial growth rates under cross-resistance and dual-resistance mechanisms. Simulated phage treatments included multispecies-targeting cocktail (1E+1S - green), pathogen-targeting cocktail of two *E. coli* phage(2E - blue), or partner-targeting cocktail of two *S. enterica* phage (2S - yellow). Arrows indicate the relative growth rates of the experimental coculture measured in panel B. The multispecies-targeting and pathogen-targeting cocktails (green and blue lines) have experimental equivalents.

We wanted to know if resistance type was always the determining factor of cocktail efficacy. Others have shown that differences in relative growth rates of cross-feeders change the recovery time from abiotic perturbations (Hom and Murray, 2014; La Sarre *et al.*, 2017; Hammarlund *et al.*, 2018). Therefore, we asked how changing relative growth rates of the cross-feeders altered phage treatment outcomes when simulating both dual- and cross-resistance mechanisms. In our experimental coculture, *E. coli* grows ~1.3x as fast as *S. enterica* (Fig 4b, Fig. 4e arrow). If we made *S. enterica* grow faster than *E. coli* (left side of fig 4e graphs), then the most effective cocktail was the pathogen-targeting cocktail (2E) or the multispecies-targeting cocktail (1E+1S), depending on the resistance type (Fig 4e). Interestingly, the more similar the relative growth rates are, the smaller the differences in efficacy of cocktail treatment strategies. This suggests that targeting the slowest growing cross-feeding partner is important for effective suppression, but predicting which cocktail suppresses the pathogen the longest depends on the evolved resistance type and relative growth rates of cross-feeders.

## Discussion

We studied the optimal way to distribute two phage among two obligate cross-feeders to best suppress one focal bacterial species. In laboratory experiments, we found that a multispecies-targeting cocktail suppressed the model pathogen, *E. coli*, longer than pathogen-targeting cocktails. The simplest explanation for this result is that pathogen-targeting cocktails are overcome by a single *E. coli* mutation which confers cross-resistance to both phage. Consistent with this, we found an evolved mucoid phenotype in pathogen-targeting cocktails which did confer cross-resistance. However, we also found that even a single *S. enterica* phage suppressed *E. coli* as well as multispecies-targeting cocktails, which cannot be explained by cross-resistance. We first hypothesized that *E. coli* evolved resistance to T7 more easily than *S. enterica* evolved resistance to P22*vir*. However, resistance to P22*vir* was more common in *S. enterica* populations than resistance to T7 was in *E. coli* populations. An alternative hypothesis was rooted in population ecology: if *S. enterica* was the rate-limiting member of the obligate cross-feeding co-culture, then reducing its population would limit growth longer than a similar reduction to *E. coli*. Experiments without phage, but where initial densities were manipulated, support this hypothesis – a low starting density of *S. enterica* causes longer suppression than a similar low starting density of *E. coli*. Subsequent modeling showed that the cause of this effect was likely the differences in growth rate: *S. enterica* grows more slowly than *E. coli*, and this slower growth interacted with the population decrease caused by phage to enhance *E. coli* suppression duration. Our results highlight a novel multispecies-targeting strategy for designing phage cocktails when pathogens obligately cross-feed with other bacteria that is affected by relative growth rates and evolved resistance type.

Our most effective cocktail strategy, the multispecies-targeting cocktail, included a phage that infected a nonpathogen, *S. enterica*, that cross-fed with our model pathogen, *E. coli*. This cocktail strategy used the ecological principle that inhibiting one cross-feeding partner effectively inhibits growth of other cross-feeding partners. By leveraging the same ecological principle, our lab previously showed that growth of a cystic fibrosis pathogen, *Pseudomonas aeruginosa*, can be inhibited by targeting its cross-feeding anaerobic partners with antibiotics (Adamowicz *et al.*, 2018). While we are not the first to consider using multispecies-targeting cocktails, others have used them with a different goal - to target co-occurring pathogens (Carson *et al.*, 2010; Lehman and Donlan, 2015; Oliveira *et al.*, 2018; Milho *et al.*, 2019; Zhao *et al.*, 2019). Additionally, others have explored phage treatment of pathogens in competitive ecological contexts, but limited their analysis to single phage treatments that targeted the focal bacterial species only (Harcombe and Bull, 2005; Brockhurst *et al.*, 2006; Wang *et al.*, 2017; Yu *et al.*, 2017; Testa *et al.*, 2019). Our research extended these foundational studies by including both pathogen-targeting and multispecies-targeting cocktails, and by addressing the role of cooperative cross-feeding between a pathogen and another coculture member. We highlight an additional way to leverage microbial ecological interactions to control pathogens.

We identified two independent factors that contributed to increased efficacy of the multispecies-targeting cocktail compared to the pathogen-targeting cocktail. First, the evolution of cross-resistance limited efficacy of the pathogen-targeting cocktail. We avoided this complication by using a multispecies-targeting cocktail strategy in which the individual phage could not infect both *E. coli* and *S. enterica*. Others have suggested alternative methods to prevent the evolution of cross-resistance. For example, Yu and colleagues designed cocktails with ‘guard’ phage that inhibit the evolution of phage resistance because they were previously experimentally evolved to infect likely-to-evolve resistant cells (Yu *et al.*, 2018). However, many researchers have described multiple rounds of phage-host coevolution suggesting that protection by guard phage may be temporary on an evolutionary time scale, although this has not been tested (Koskella and Brockhurst, 2014; Jariah and Hakim, 2019). Others have used molecular techniques to identify phage binding sites and subsequently design cocktails that use multiple binding sites to increase both the number of mutations required for resistance and the cost of resistance (Filippov *et al.*, 2011). While this would protect against receptor-mediated evolution of resistance, it would not prevent general resistance mechanisms that inhibit phage access to the cell surface, such as the evolution of mucoidy, which we observed when treating the cocultures with the pathogen-targeting cocktail. Yet, others have described phage that degrade this mucoid barrier and facilitate infection by other phage (Kim *et al.*, 2015). We identified an additional method for preventing cross-resistance from reducing the efficacy of phage cocktails.

Second, we found that including a phage that targeted the slower-growing cross-feeding partner was key to effectively suppressing pathogen growth. We used mathematical simulations to determine that relative growth rates of the cross-feeding partners altered how effective including a phage targeting the slower-grower was (Fig 4e). In fact, inhibiting the slower-grower, *S. enterica*, with a single phage was as effective as inhibiting with the multispecies-targeting cocktails in experiments (Fig 1b) and in simulations (Fig 4d). Our findings agree with other studies that suggest that changes in relative growth rates of community members (Banks and Bryers, 1991; Raskin *et al.*, 1996), particularly cross-feeders (Turner *et al.*, 1996; Hammarlund *et al.*, 2018) can alter responses to perturbations. To expand on these foundational studies, we used mathematical simulations to explore how relative growth rates impact the magnitude of response to perturbations. We found that the more similar the relative growth rates of the cross-feeders were, the smaller the difference in efficacy of cocktail strategies. Conversely, the more different the relative growth rates were, the more benefit we observed in targeting the slower-growing cross-feeding partner. While our simulations suggest that a *S. enterica*-targeting cocktail would be most effective at suppressing *E. coli* if cross-resistance did not evolve (Fig 4e), we were unable to test this because our efforts to find a second *S. enterica* phage that replicated in our coculture were unsuccessful (Fig S1). Our results suggest that including at least one phage targeting the slower-grower in a cross-feeding coculture is an effective method to extend pathogen suppression. Furthermore, our results indicate that if the relative growth rates of a pathogen and its cross-feeding partner are unknown, adding a nonpathogen-targeting phage could be one way to maximize the odds of inhibiting the pathogen.

A complication in a clinical setting or agricultural application could be that absolute pathogen population size, or pathogen load, may be more critical to treatment outcomes than how long the growth of a pathogen can be suppressed. Here, in two of fifteen communities treated with the T7 *E. coli*-targeting phage either alone or in a cocktail we observed complete eradication of *E. coli* populations and lower final *E. coli* densities in cocultures in which *E. coli* was not eradicated (Fig S3). This would indicate that directly targeting the pathogen would be the fastest way to immediately decrease pathogen load. But the rate of resistance evolution would determine how long population sizes are kept low. Our results indicate that including phage targeting cross-feeding partners is one way to limit the recovery of knocked-down pathogen populations. These approaches are not mutually exclusive - phage cocktails could include multiple phage targeting the pathogen, and one or more phage targeting its cross-feeding partner.

An alternative method for targeting multiple species in a community is with polyvalent phage treatment, or phage with host ranges that encompass multiple species. Descriptions of polyvalent phage have increased over the past five years likely due to directed changes in phage isolation protocols (Hamdi *et al.*, 2017; Duc *et al.*, 2020; Li *et al.*, 2020). In fact, Zhao and colleagues used a soil-carrot microcosm system to compare the efficacy of a cocktail that included phage targeting two different plant pathogens with a treatment of a single polyvalent phage that infected both pathogens (Zhao *et al.*, 2019). They found that both treatments effectively limited the growth of both pathogens, but the polyvalent phage treatment disturbed the soil microbiome less than the multipathogen-targeting cocktail. Some challenges with using polyvalent phage might include differences in host preference based on receptor-phage binding strength. If binding strength were different enough, the polyvalent phage should function like a phage that targeted a single species. However, one benefit is that phage populations could grow faster because more hosts would be available, although no studies have directly tested this yet. We suggest that future research could test including polyvalent phage with different cocktail strategies.

In conclusion, we have illustrated a novel phage cocktail strategy for targeting cross-feeding pathogens. Our strategy limits cross-resistance evolution and maximizes pathogen suppression by targeting both the slower-growing partner and the pathogen. These and other results indicate that leveraging microbial community ecological interactions is a promising approach to help control pathogen growth in a variety of applications in human health, agriculture, and food safety.

## Experimental Procedures

### Bacterial and Phage Strains in the Cooperative Co-Culture System

The bacterial strains used in this experiment have been previously described (Fig 1A)(Harcombe, 2010). Briefly, the *E. coli* K12-derivative has a *metB* deletion and cyan fluorescent protein (CFP) in the attB lambda integration site. *S. enterica* is an LT2 strain with mutations in *metA* and *metJ* causing methionine secretion and yellow fluorescent protein (YFP) in the *attB* lambda integration site (Douglas *et al.*, 2016, 2017). *E. coli* metabolizes lactose and excretes acetate which *S. enterica* consumes. *S. enterica* excretes methionine which is used by *E. coli*. Bacterial stocks were stored at −80°C in 20% glycerol. *E. coli*-specific phage T7 and P22*vir* were provided by Ian Molineaux (UT Austin) and *S. enterica*-specific P1*vir* by Ross Carlson (Montana State University). Phage stocks were grown on monocultures of ancestral *E. coli* or *S. enterica* in lactose or acetate minimal medium at 30°C. Cells were lysed with chloroform, centrifuged to pellet cell debris, and stored at 4°C.

Monoculture and co-culture experiments used a defined minimal medium (14.5 mM K_2_HPO_4_, 16.3 mM NaH_2_PO_4_, 0.814 mM MgSO_4_, 3.78 mM Na_2_SO_4_, 3.78 mM (NH_4_)_2_SO_4_) supplemented with trace metals (1.2 μM ZnSO_4_, 1 μM MnCl_2_, 18 μM FeSO_4_, 2 μM (NH_4_)_6_Mo_7_O_24_, 1 μM CuSO_4_, 2 mM CoCl_2_, 0.33 μM Na_2_WO_4_, 20 μM CaCl_2_) as described (Delaney *et al.*, 2013). Carbon sources were 2.78 mM D-lactose or acetate, as indicated. Monocultures of *E. coli* were supplemented with 20μM L-methionine.

### Measuring E. coli Suppression in the Cross-Feeding Co-Culture

Bacterial growth at 30°C was tracked every 20 min with OD600 and fluorescence measurements using a shaking plate reader (Tecan Infinite ProM200). *E. coli* was measured with CFP (Ex: 430 nm; Em: 490 nm), and *S. enterica* with YFP (Ex: 500 nm; Em: 530 nm). We used four - five replicates of each treatment, as indicated. To wells in a 96-well plate, 10^5^ cells each of mid-log phase *E. coli* and *S. enterica* were inoculated into 200μL of lactose minimal media with 5×10^2^ virions as indicated (MOI = 0.05 per phage). Cultures incubated for 5 days until stationary phase was reached. *E. coli* suppression length in hours was estimated by calculating the time to 95% maximum CFP measurement.

### Profiling Resistance to Phage via Cross-Streak Assays

To assay for acquired phage resistance, we used cross-streak assays with representative isolates from treatments. 30μL of ancestral phage stock (10^8^ to 10^9^ PFU/mL) was dripped down a minimal medium agar plate and left to dry. Overnight cultures of isolates grown in minimal medium were streaked perpendicular to the phage. Plates were incubated at 30°C until growth was visible. Isolates were determined to be resistant if streaks were uniform across the phage line and sensitive if bacterial growth was interrupted.

### Resistance to Phage due to Standing Variation in Ancestral Bacterial Stocks

To determine frequency of phage resistance of ancestral bacteria, we quantified the number of cells that grew on phage-saturated agar plates. Ancestral *E. coli* and *S. enterica* monocultures were grown in lactose + methionine or acetate minimal media, respectively, for 3 days at 30°C. LB plates were saturated with 1ml of ancestral phage stock (~1×10^9^ PFU/mL), dried, and spotted with 5μl of bacterial monocultures in 10-fold dilutions. Plates were incubated at 30°C until phage-resistant colonies were counted. We compared the number of colonies of plates with and without phage for each phage-host combination.

### Assessing the Effect of Starting Frequency of Microbial Partners on Coculture Growth

We tested the time to maximum density of *E. coli* in the coculture when starting frequencies were altered in the absence of phage We started the rare partner of cocultures at 0.03% while holding the common species at 10^5^ cells/well in lactose minimal medium (n=5). Community growth was as described above (see Methods: *Measuring Experimental Cross-Feeding…).*

### In silico Modeling of Communities

To represent our cross-feeding microbial community, we modified a series of resource-explicit ordinary differential equations to simulate an *E. coli* and *S. enterica* cross-feeding system in which one species grows on nutrients secreted by the other (Fazzino *et al.*, 2020). We used Monod equations with multiplicative limitation of lactose and methionine essential nutrients for *E. coli*. The model mimics the metabolic network of the synthetic experimental coculture.

The major metabolites – lactose, acetate, and methionine – are tracked throughout simulations. Lactose is seeded in and is depleted as *E. coli* grows. Acetate is produced by *E. coli* growth, and is depleted by *S. enterica* growth. Methionine is produced during *S. enterica* growth, and is depleted during *E. coli* growth. Simulated cocultures grow until all lactose is consumed.

Each species has multiple genotypes to simulate resistance to different phage, with the amount seeded in representing mutation rarity. Resistant genotypes had founder population sizes at a maximum of 0.1% of the sensitive genotype to simulate rare resistance. *E. coli* had four genotypes; Es for sensitive to both phage, ErT7 for resistant to only T7, ErP1 for resistant to only P1*vir*, and ErT7P1 for resistant to both phage. *S. enterica* had two genotypes; Ss for sensitive to P22*vir*, and Sr for resistant to P22*vir*. Resistance was modeled as complete and without cost. The replication of each phage strain– T7, P1*vir*, and P22*vir* – was determined by adsorption rates and burst sizes. Each phage species can only kill sensitive genotypes of a single bacterial species. Model parameters are informed by literature values and were parameterized to approximate coculture growth dynamics without phage (Fig 4, Table S2)(Fazzino *et al.*, 2020). Bacterial growth rates were measured from wet-lab monoculture experiments (*E. coli* grown in lactose + methionine and *S. enterica* grown in acetate) where OD600 was measured every 20min. Growth curves were fit with a non-linear least-squares Baranyi function of the growth rate parameter, as described (Baranyi and Roberts, 1994).

Figure 4 shows a schematic representation of the following equations and parameters.

*E. coli* (E) growth:

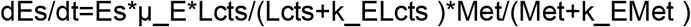

*S. enterica* (S) growth:

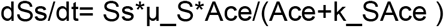

Population sizes (E or S) are multiplied by their species-specific growth rates per hour (μ_x_) and a Monod saturation function with a species and resource explicit constant (k) for each necessary resource. During phage infection, cell lysis is simulated. For example, when P1vir infects an *E. coli* that is only T7-resistant:

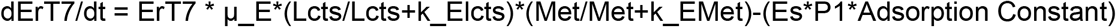

New phage are added with host death:

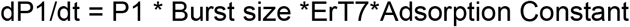

*E. coli*, *S. enterica*, and phage equations are repeated for each individual genotype and phage.

Simulations were run in R with the DeSolver package, using the LSODA solver (Soetaert *et al.*, 2010).

## Acknowledgements

We would like to thank Sarah P. Hammarlund, Harcombe Lab members, and the UMN Institute for Molecular Virology community for useful discussions. L. Fazzino was supported by a Fellowship from the Institute for Molecular Virology Training Program (NIH T32 AI083196). Research was also supported through the NIH (GM121498-01A1, to William Harcombe). The authors declare no conflict of interest.

