## Supplemental for "Phage cocktail strategies for the suppression of a pathogen in a cross-feeding coculture"

### Supporting Figures & Tables

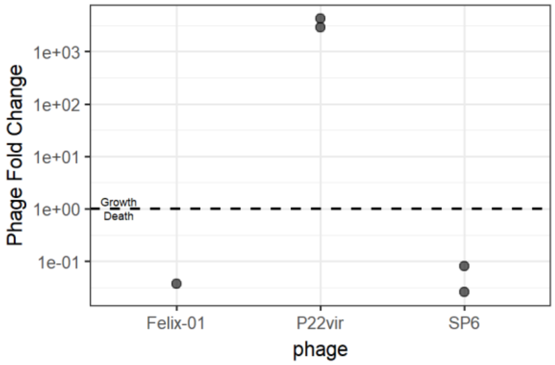


**Supplemental Figure 1. Screening of *S. enterica*-specific phage activity in cooperative coculture.** P22*vir*, SP6, and Felix-01 *S. enterica*-specific phages were inoculated into *E. coli*-*S. enterica* cocultures and grown at 30°C while shaking until stationary phase was reached (4-5 days, n = 1-2). Initial and final PFU/ml were measured by plating with ancestral *S. enterica.* Only P22*vir* increased in concentration over the growth period.

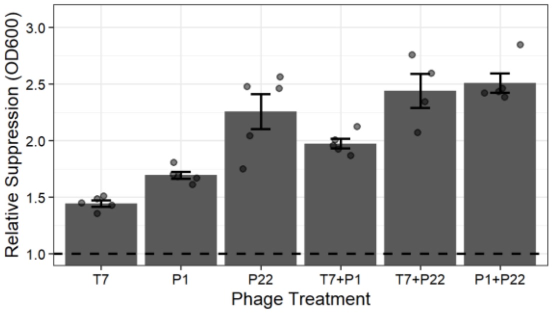


**Supplemental Figure 2. Coculture-level suppression lengths caused by phage treatments.** Relative coculture suppression lengths of single and cocktail phage treatments standardized to the no phage control. Suppression length was calculated using 95% maximum OD600. Bars represent means ± SE (n = 4-5).

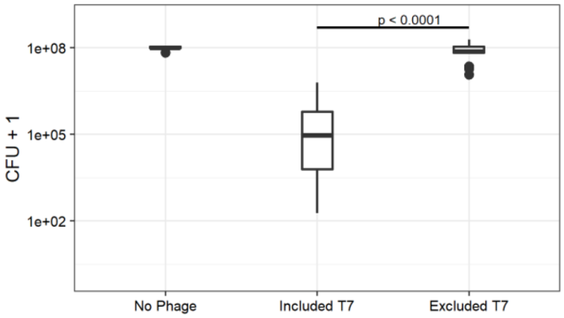


**Supplemental Figure 3. Boxplots of final *E. coli* densities after phage treatments.** Including T7 phage in treatments lowered final *E. coli* population size. Cocultures were grown with single phage treatments and cocktails and bacterial populations sizes were counted by plating with selective plates. Statistical significance was tested with a Two-sample Mann–Whitney U. (n = 15)

**Table S1. Absolute and relative suppression lengths of phage treatments.**

| Phage Treatment | Treatment Type | Absolute Suppression Length (hrs ± SE) | Relative Suppression Length (standardized to phage-free) |
|---|---|---|---|
| Phage-free | None | 34.44 ± 0.0 | 1.00 ± 0.0 |
| T7 | Single phage | 49.74 ± 2.05 | 1.44 ± 0.06 |
| P1vir | Single phage | 58.37 ± 2.35 | 1.69 ± 0.07 |
| P22vir | Single phage | 77.7 ± 12.0 | 2.26 ± 0.35 |
| T7+P1vir | Pathogen-targeting | 67.98 ± 3.28 | 1.97 ± 0.10 |
| T7+P22vir | Multispecies-targeting | 84.03 ± 10.42 | 2.44 ± 0.30 |
| P1vir + P22vir | Multispecies-targeting | 86.3 ± 6.56 | 2.5 ± 0.19 |

**Table S2. Parameters for resource-explicit ODE mathematical model.**

| Parameter (name in model) | Parameter value |
|---|---|
| *E. coli* growth rate (mu_e) | 0.291/hr |
| *S. enterica* growth rate (mu_s) | 0.221/hr |
| *E. coli* production of ace (p_e_ace) | 4e-12 grams produced/*E. coli* cell |
| *S. enterica* consumption of ace (c_s_ace) | 3e-12 grams consumed/*S. enterica* cell |
| *S. enterica* production of met (p_s_met) | 4e-12 grams produced/*S. enterica* cell |
| *E. coli* consumption of met (c_s_met) | 3e-12 grams consumed/*E. coli* cell |
| *E. coli*-specific T7 burst size (burst_T7) | 100 phage/burst *E. coli* cell |
| *E. coli*-specific T7 adsorption rate | 1e-9 /phage*E. coli* cell |
| *E. coli*-specific P1*vir* burst size (burst_P1) | 100 phage/burst *E. coli* cell |
| *E. coli*-specific P1*vir* adsorption rate | 1e-9 /phage*E. coli* cell |
| *S. enterica*-specific P22*vir* burst size (burst_P22) | 100 phage/burst *S. enterica* cell |
| *S. enterica*-specific P22*vir* adsorption rate | 1e-9 /phage*E. coli* cell |

**Table S3. Starting densities of cross- and dual-resistance modeling.**

| Simulated Organism | Organism Description | Cross-Resistance | Dual-Resistance |
|---|---|---|---|
| Es | Sensitive *E. coli* | 9.98999e5 | 9.97999e5 |
| ErT7 | T7-resistance *E.coli* | 0 | 1e3 |
| ErP1vir | P1vir-resistant *E. coli* | 0 | 1e3 |
| Er | T7- and P1vir-resistant *E. coli* | 1.001e3 | 1 |
| Ss | Sensitive *S. enterica* | 9.98999e5 | 9.97999e5 |
| SrP22vir | P22vir-resistant *S. enterica* | 0 | 1e3 |
| Sr2 | Phage2-resistant *S. enterica* [a] | 0 | 1e3 |
| Sr | P22vir and Phage2-resistant *S. enterica* | 1.001e3 | 1 |
| Total Biomass | Sum of all bacterial biomass | 1e6 | 1e6 |

[a] Phage 2 is for simulation purposes only and does not correspond to a second experimental *S. enterica* phage.
